# SCL28 promotes cell expansion and endoreplication in Arabidopsis by activating *SIAMESE-RELATED* cyclin-dependent kinase inhibitors

**DOI:** 10.1101/2021.08.10.455231

**Authors:** Camila Goldy, Virginia Barrera, Isaiah Taylor, Celeste Buchensky, Rodrigo Vena, Philip N. Benfey, Lieven De Veylder, Ramiro E. Rodriguez

## Abstract

The processes that contribute to plant organ morphogenesis are spatial-temporally organized. Within the meristem the mitotic cell cycle produces new cells that subsequently engage in specific cell expansion and differentiation programs once they exit the division competent zone. The latter is frequently accompanied by endoreplication, being an alternative cell cycle that replicates the DNA without nuclear division, causing a stepwise increase in somatic ploidy. We have previously shown that the Arabidopsis SCL28 transcription factor promotes progression through G2/M and modulates division plane orientation. Here, we demonstrate that *SCL28* co-express and regulates genes specific to cell elongation and differentiation, including genes related to cell wall and cytoskeleton assembly. Consistently, this correlates with defects in post-mitotic cell expansion in a *scl28* mutant. Strikingly, SCL28 controls expression of 6 members of the *SIAMESE/SIAMESE-RELATED* (*SIM/SMR*) family, encoding cyclin-dependent kinase inhibitors with a role in promoting mitotic cell cycle exit and endoreplication onset, both in response to developmental and environmental cues. Consistent with this role, *scl28* mutants displayed reduced endoreplication, both in roots and leaves. Altogether, these results suggest that *SCL28* controls cell expansion and differentiation by promoting endoreplication onset and by modulating aspects of the biogenesis, assembly and remodeling of the cytoskeleton and cell wall.

## Introduction

After new cell types are generated by stem cells in plant meristems, cells go through a stage of active proliferation in which small and generally isodiametric cells with limited differentiation are produced through the mitotic cell cycle (MCC). Later, anisotropic cell expansion and cell differentiation proceeds until cells reach their mature functional state. In every plant organ, each of these processes takes place in clearly defined regions (Heidstra and Sabatini, 2014). For example, towards the tip of a growing root, cells actively divide in the meristematic zone. Shootward, cells exit the mitotic cell cycle in the transition zone and later, in the elongation and differentiation zones cells increase in size and differentiate (Petricka et al., 2012). Local parameters of cell division and expansion, such as the number of cells produced in meristems, the orientation of cell division and the direction of anisotropic cell expansion are important in determining the final size and shape of plant organs.

For cells to elongate in a particular direction during post mitotic anisotropic cell expansion, cell walls must undergo specific and local chemical modifications. Local softening or stiffness of the cell wall is essential to change from isotropic to anisotropic growth, which in turn is crucial to determine cell size and shape (Wolf et al., 2012). The plant cell wall is a complex network composed of rigid cellulose microfibrils embedded in a matrix of hemicellulose, pectins, proteins and other biomolecules. The mechanical properties of the cell wall are highly influenced by the chemical state of its components. For example, changes in pectin chemistry, such as deesterification or deacetylation, correlate with changes in cell wall elasticity (Braybrook and Peaucelle, 2013; Haas et al., 2021). Also, modifications of the structure of hemicelluloses modulate the properties of the cell wall and, therefore, influence plant development (Stratilova et al., 2020). The cytoskeleton plays important roles in plant cell and tissue morphogenesis. Cortical microtubules, which organize in a region underlying the plasma membrane are highly dynamic structures and are crucial in determining cell growth orientation (Chebli et al., 2021).

In the transition zone, cells exit MCC, but frequently perform some degree of endoreplication (ER) (Bhosale et al., 2018) (Hayashi et al., 2013). During this alternative cell cycle, also known as endocycle, DNA is replicated without nuclear or cytoplasmic division causing a stepwise increase in nuclear DNA content (Bhosale et al., 2019; Lang and Schnittger, 2020). Endoreplication is involved in several plant development pathways and has been correlated with post mitotic cell growth, cell differentiation, high metabolic activities, rapid anisotropic cell expansion, and with the ability to respond to DNA damage (Robinson et al., 2018; Bhosale et al., 2019; Tsukaya, 2019). Remarkably, different cell types of a plant organ perform characteristic rounds of ER cycles reaching specific somatic polyploidy levels (Katagiri et al., 2016; Bhosale et al., 2018). For example, in the root vascular bundle, xylem cells reach high levels of ploidy while those of the phloem remain almost exclusively diploid (Bhosale et al., 2018). This indicates that the transition from the MCC to endoreplication and the magnitude of the endoreplication itself are precisely spatially and temporally regulated.

The mechanisms that promote the transition from the MCC to endoreplication involve a variety of pathways that specifically suppress the activity of mitotic cyclin/cyclin-dependent kinases complexes (CYC-CDK) leaving the replication machinery functional with cells alternating between S and G1 phases. One mechanism involves the degradation of mitotic CYC-CDK complexes by the Anaphase/Cyclosome Promoting Complex (APC/C). This complex, assisted by CCS52A activators, acts as a Ubiquitin Ligase E3 complex that marks rate-limiting cell cycle proteins for destruction. Two different CCS52A proteins, CCS52A1 and CCS52A2, are encoded in the Arabidopsis genome (Cebolla et al., 1999; Tarayre et al., 2004). Both APC/C regulators promote endoreplication onset, as evidenced by reduced DNA ploidy levels in mutant plants (Lammens et al., 2008; Kasili et al., 2010; Heyman et al., 2017).

Another pathway involves the association of CDKs with inhibitory proteins of the *SIAMESE/SIAMESE-RELATED* (*SIM*/*SMR*) family. *SIAMESE*, the founding member of this family of cyclin-dependent kinase inhibitors (CKI), was initially discovered through its function in trichome development and endoreplication control (Churchman et al., 2006). There are 17 *SIM/SMR* genes in the Arabidopsis genome (Yi et al., 2014; Kumar et al., 2015). Among the characterized *SMRs*, *SIM*, *SMR1/LGO* and *SMR2* promote MCC exit and endoreplication onset during leaf and root development (Churchman et al., 2006; Kasili et al., 2010; Roeder et al., 2010; Bhosale et al., 2018), *SMR5* and *SMR7* control cell cycle arrest in response to genotoxic stress (Yi et al., 2014; Bourbousse et al., 2018; Takahashi et al., 2019), whereas *SMR4* and *SMR8* promote the transition from proliferation to differentiation within the stomatal lineage (Han et al., 2021).

We have previously shown that in Arabidopsis, SCL28, a GRAS transcription factor, promotes progression through G2/M in mitotic cells and modulates the selection of cell division planes, phragmoplast activity and mitotic cell expansion. Here, we extend the functional characterization of SCL28 in plant development, revealing a previously unrecognized role for SCL28 in post-mitotic cell expansion and endoreplication onset through the modulation of the expression of a subset of *SIM/SMRs*, cytoskeleton and cell wall genes.

## Results

### SCL28 promotes organ growth by modulating cell expansion

We have previously shown that *SCL28* promotes root growth (Goldy et al., 2021) (Figure S1A-C). We extended the characterization of the function of *SCL28* by analyzing shoot phenotypes of plants carrying an insertional mutant allele (*scl28-3*) in which *SCL28* mRNA is almost undetectable (Figure S2). We found that in the mutant, rosettes are smaller and inflorescence stems shorter (Figure S1D, E, G and H) due to a reduction in rosette and stem growth rate (Figure S1F). These growth defects were complemented by transforming *scl28-3* with a construct including the *SCL28* coding sequence fused to the fluorescent protein VENUS under the control of *SCL28* upstream regulatory regions (*ProSCL28:SCL28-VENUS*) (Figure S1 and S2).

As organ growth results from the combined action of cell proliferation and expansion, we evaluated mature cell size in multiple organs. We found that *scl28-3* plants display a generalized defect in cell expansion. Cells in the root cortex and epidermis, hypocotyl epidermis, inflorescence stem epidermis, leaf palisade parenchyma and leaf epidermal cells, were on average 30% smaller in the mutant as compared to wild type (Figure 1 and Table S1). These defects in cell expansion were complemented when *scl28-3* was transformed with the *ProSCL28:SCL28-VENUS* construct. Also, when we analyzed plants transformed with a repressor version of *SCL28* (*ProSCL28:SCL28-SRDX*) (Goldy et al., 2021) we found similar phenotypes to those observed in *scl28-3*, including a reduction in root growth rate (Figure S3A and B) and defects in root and leaf cell expansion (Figure S3C-J). Finally, overexpression of *SCL28* in β-estradiol-treated *XVE-SCL28* plants (Figure S5A) resulted in a moderate increase in root growth rate (Figure S4A-B) accompanied with an average increase of 20% in the size of root and leaf cells (Figure S4C-J). Taken together, these results indicate that *SCL28* promotes both shoot and root growth by modulating not only cell proliferation but also cell expansion.

**Figure 1.**
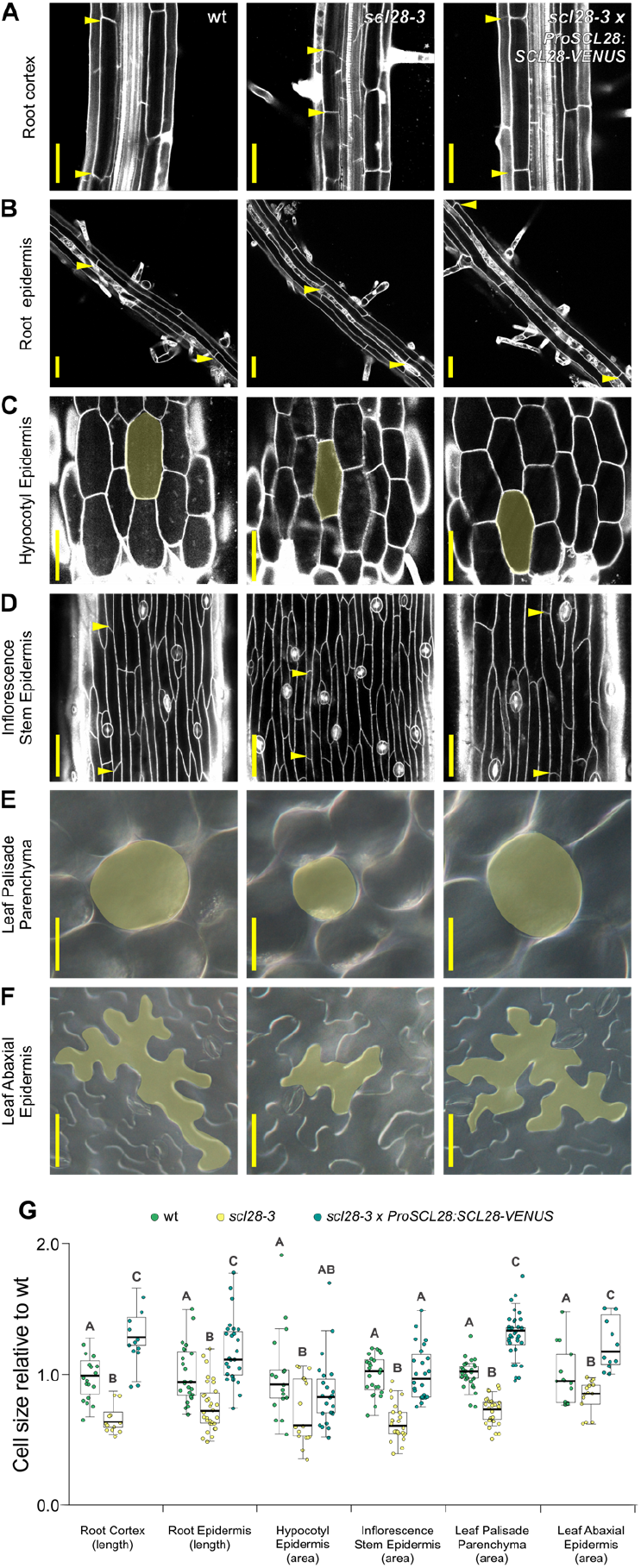
SCL28 promotes cell expansion. **A-F.** Micrographs showing cells from root cortex (A), root epidermis (B), hypocotyl epidermis (C), inflorescence stem epidermis (D), leaf palisade parenchyma (E) and leaf abaxial epidermis (F) in wild-type (wt), *scl28-3* and *scl28-3* × *ProSCL28:SCL28-VENUS* plants. Images from panels A to D were obtained by LSCM in propidium iodide (PI) stained (grayscale) organs. Images from panel E and F were obtained by DIC microscopy in fixed and cleared leaves. Scale bars, 50 μm. The limits of representative cells are labelled with arrow heads (A, B, and D) or shaded in yellow (C, E and F). **G.** Cell size measurements normalized to the average value obtained in wild-type (wt) plants. Different letters indicate significant differences (*P* < 0.05; ANOVA followed by Tukey’s multiple comparison test).

### Transcriptome analysis of scl28-3 reveals roles for SCL28 in the transition and elongation zones

To define the networks regulated by *SCL28*, we performed transcriptome profiling of wild-type and *scl28-3* root tips. We found 777 differentially expressed genes (DEGs), with 341 up-regulated and 436 down-regulated in *scl28-3* compared to wild type (*p* < 0.05 and fold change > 25%; Table S2).

We first analyzed the expression pattern of these DEGs in a root developmental stage-specific gene expression dataset (Figure 2A) (Wendrich et al., 2017). As expected from the already described functions of *SCL28*, ~50% of the DEGs detected in the dataset are enriched in the proximal region of the root meristem where cell proliferation occurs at the highest rate and where mitotic markers are preferentially induced (Figure 2B and C). The other ~50% of the DEGs detected in the dataset are enriched in the medial and distal zones, where genes related to endoreplication, cell wall organization and biogenesis and cell differentiation are enriched (Figure 2B and C) (Wendrich et al., 2017). These results suggest that *SCL28* might have functions outside the cell proliferation domain and are consistent with the defects in cell expansion described above. Accordingly, the gene expression database indicated that *SCL28* mRNA is detected across all root developmental zones in a gradient with a peak in the meristematic zone declining shootward (Figure 2A) (Wendrich et al., 2017). Expression of the *ProSCL28:SCL28-VENUS* reporter confirmed this expression pattern with highest levels in the root meristem (Goldy et al., 2021) and reduced, but detectable, expression in the transition and elongation zones (Figure 2D).

**Figure 2.**
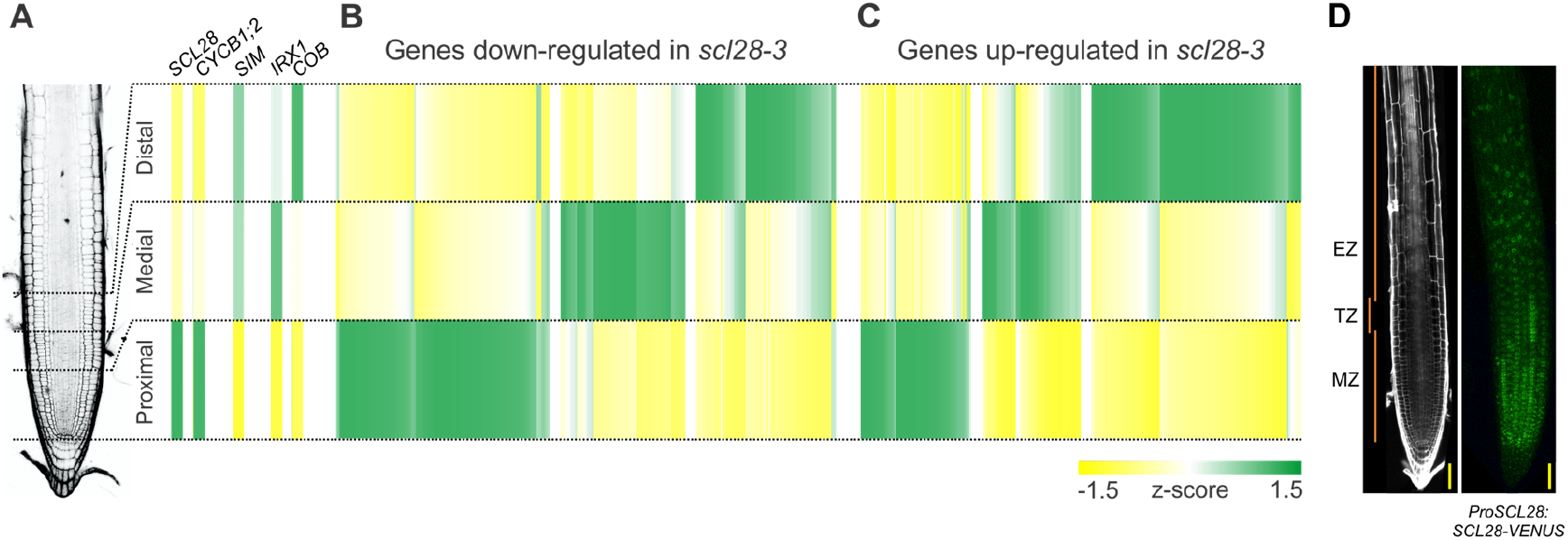
SCL28 modulates the expression of genes across all developmental zones of the root. **A.** Expression along the root’s longitudinal axis of *SCL28, CYCB1;2* as a mitotic marker, *SIAMESE* (*SIM*) as an endoreplication marker and *IRREGULAR XYLEM 1* (*IRX1*) and *COBRA* (*COB*) as differentiation markers. **B-C.** Expression along the root’s longitudinal axis of genes down-regulated (B) or up-regulated (C) in *scl28-*3 roots. For panels A to C, data was obtained from the literature (Wendrich et al. (2017) PNAS), z-scores were calculated for the detected genes in the dataset and expressed as a heat map. This data set includes 3 regions of the root named by the authors as proximal, medial, and distal. **D.** Expression pattern of *SCL28* determined by LSCM with a *ProSCL28:SCL28-VENUS* reporter in 6-d-old roots. The image is a maximum intensity projection of a z-stack obtained from the epidermis to the medial longitudinal section of the root. Scale bars, 50 μm. MZ, meristematic zone; TZ, transition zone; EZ, elongation zone.

As *scl28-3* displays defects in cell expansion, we inspected the DEG lists for genes related to cytoskeleton and cell wall biogenesis (Table S2). Among these, we found that *IQ-DOMAIN 16/ABNORMAL SHOOT 6* (*IQD16/ABS6*) is down regulated in *scl28-3*. *IQD16/ABS6*, together with *KATANIN1* and *SPIRAL2*, have been shown to co-localize with microtubules and participate in microtubule spatial organization during mitosis and anisotropic cell expansion (Burstenbinder et al., 2017; Wendrich et al., 2018; Yang et al., 2020; Li et al., 2021)

Another gene down regulated in *scl28-3*, *ACTIN DEPOLYMERIZING FACTOR 6* (*ADF6*) is a member of the actin-depolymerizing factors (ADF) family of actin binding proteins involved in generating and remodeling the actin cytoskeleton (Ruzicka et al., 2007). We also found several genes related to cell wall biogenesis, assembly, and remodeling, including expansins, extensins, hydrolases, extracellular peroxidases, pectin methyl esterases, endoxyloglucan transferases and cell wall structural proteins (Table S2). Taken together, these results suggest that *SCL28* could control cell expansion and differentiation by modulating aspects of the biogenesis, assembly and remodeling of the cytoskeleton and cell walls.

### SCL28 modulates expression of genes related to cytoskeleton and cell wall assembly

To validate the results obtained with the transcriptome analysis we used RT-qPCR to determine the expression of a subset of genes related to cytoskeleton and cell wall assembly (Figure 3A). We found that *IQD16*, *ADF6* and several cell wall assembly genes displayed changes in expression consistent with those observed in the transcriptome analysis. When *SCL28* was overexpressed in *XVE-SCL28* plants by a short treatment with β-estradiol that does not generate evident phenotypes (Figure S5), all these genes displayed opposite changes in expression (Figure 3A), supporting the idea that the GRAS TF might regulate their expression.

**Figure 3.**
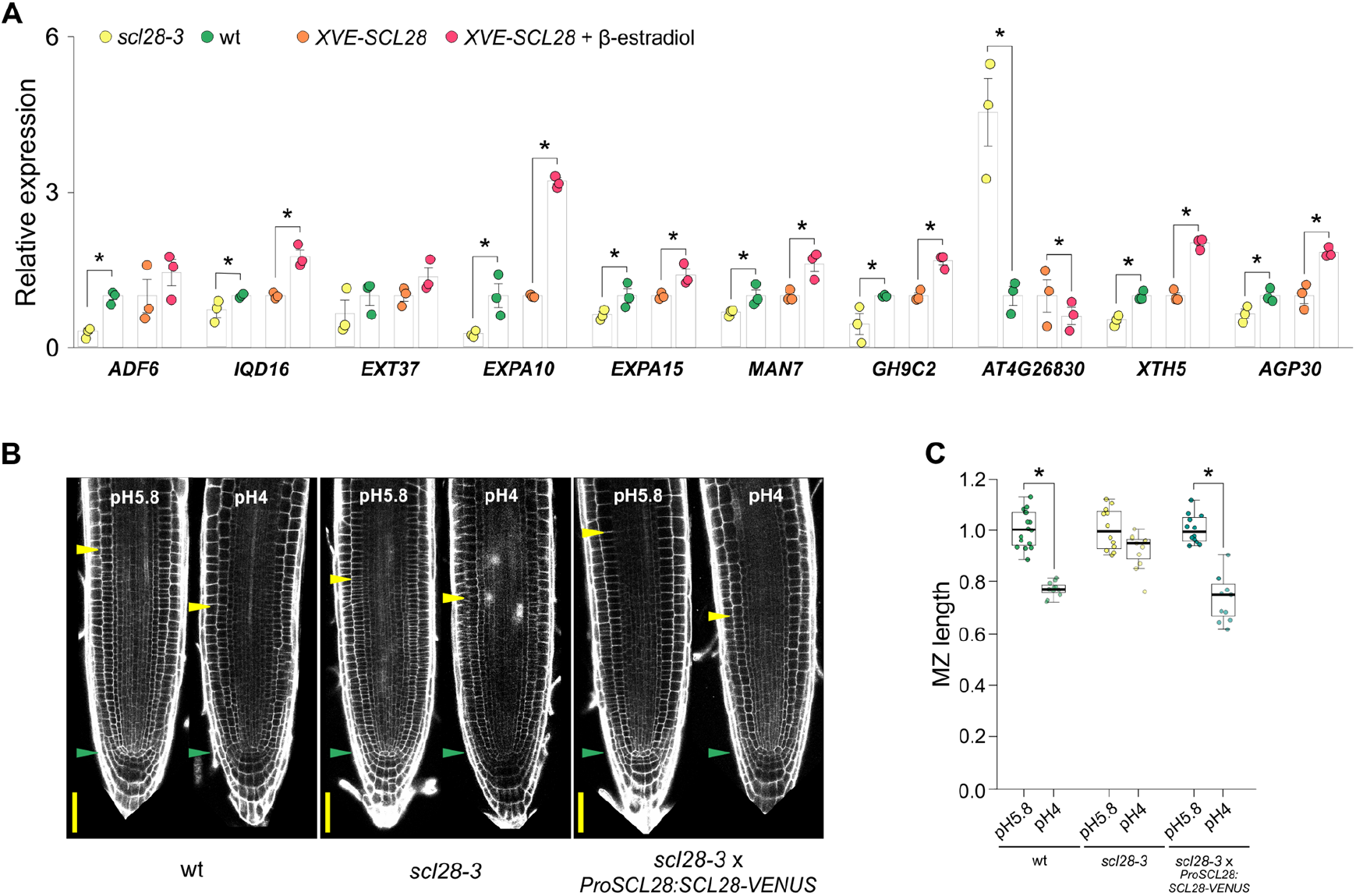
SCL28 modulates the expression of genes related to cytoeskeleton and cell wall assembly. **A.** Expression of genes related to cytoskeleton (*ADF6* and *IQD16*) and cell wall (*EXT37*, *EXPA10*, *EXPA15*, *MAN7*, *GH9C2*, *AT4G26830*, *XTH5* and *AGP30*) assembly in wild-type (wt), *scl28-3* and *XVE-SCL28* plants, the latter incubated for 16 h in MS media supplemented with 0.25 μM β-estradiol. Expression was determined by RT-qPCR in three biological replicates and normalized to the mean value obtained in wild-type plants. Asterisks indicate significant differences (*P* < 0.05, Student’s t test comparing wild type with scl28-3 or mock vs β-estradiol-treated *XVE-SCL28* plants). **B.** Root tip architecture of 6-d-old wild-type (wt), *scl28-3* and *scl28-3* × *ProSCL28:SCL28-VENUS* plants grown on standard (pH 5.8) or acidic (pH 4.0) conditions. Images were obtained by LSCM in PI (grayscale) stained plants. Green and yellow arrowheads mark the position of the QC and the end of the meristem where cells start to elongate. Scale bars, 50 μm. **C.** Meristematic zone length of 6-d-old wild-type (wt), *scl28-3* and *scl28-3* × *ProSCL28:SCL28-VENUS* plants grown on standard (pH 5.8) or acidic (pH 4.0) conditions. Values were normalized to the average value obtained in standard pH within each genotype. Asterisks indicate significant differences between growth conditions (*P* < 0.05, Student’s t test).

Among the genes activated by *SCL28, EXPANSIN A10* (*EXPA10)* and *EXPANSIN A15* (*EXPA15)* code for cell-wall located α-expansins that promote cell expansion when activated by low apoplastic pH (Cosgrove, 2005). It has been shown that these expansins define the position of the transition zone by initiating cell expansion when activated by acidification of the apoplast (Pacifici et al., 2018). Consistent with this, while mild acidification of the growth media reduced the size of the root meristem in wild-type plants (Figure 3B), no effect on root meristem size was observed in *scl28-3* (Figure 3B), presumably due to the low expression levels of *EXPA15* and *EXPA10* found in the mutant (Figure 3A). Importantly, the sensitivity to low pH was restored when the *ProSCL28:SCL28-VENUS* construct was introduced into *scl28-3* plants (Figure 3B).

### SCL28 promotes endoreplication and cell expansion by activating SMR13

Gene Ontology (GO) enrichment analyses showed that the list of genes down-regulated in *scl28-3* is enriched in genes related to DNA replication and endoreplication (Table S2). In particular, 6 of the 17 members of the *SIM/SMR* family (Yi et al., 2014; Kumar et al., 2015) were down regulated in *scl28-3* (Table S2), suggesting a role for SCL28 in promoting MCC exit and endoreplication onset. Consistent with SCL28 controlling the expression of these genes, the mRNA levels of *SMR2*, *−6*, *−7*, *−9* and *−13* correlated with *SCL28* mRNA levels when measured by qPCR in *scl28-3* or β-estradiol*-*induced *XVE-SCL28* plants (Figure 4A). No significant changes were detected for *SIM* or *SMR1* mRNAs (Figure 4A). Also, no reduction in expression was detected in *scl28-*3 for genes coding for CKIs from the *ICK/KRP* gene family (Figure S6). Finally, *CCS52A1* or *CCS52A2*, which code for APC/C activators that promote endoreplication onset showed no significant reduction in expression in *scl28-3* (Figure S6). These results indicate that SCL28 activates the expression of a specific subset of cyclin-dependent kinase inhibitors belonging to the *SIM/SMR* family.

**Figure 4.**
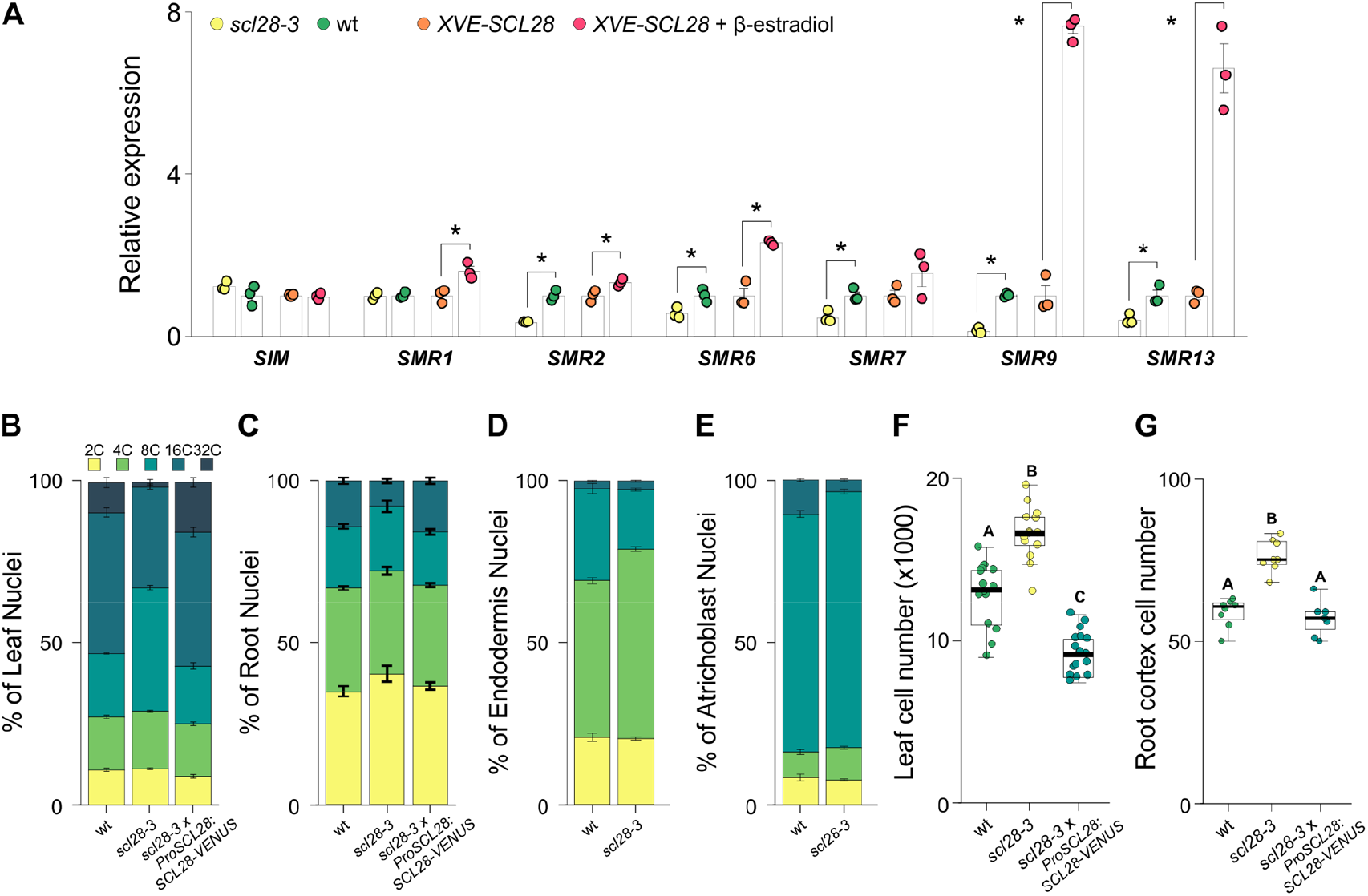
SCL28 promotes endoreplication onset by activating the expression of SIAMESE-RELATED cyclin-dependent kinase inhibitors. **A.** Expression of *SIAMESE-RELATED* genes in wild-type (wt), *scl28-3* and *XVE-SCL28* plants, the latter incubated for 16 h in MS media supplemented with 0.25 μM β-estradiol. Expression was estimated by RT-qPCR in three biological replicates and normalized to the mean value obtained in wild-type plants. Asterisks indicate significant differences (*P* < 0.05, Student’s t test comparing wild type with *scl28-3* or mock vs β-estradiol treated *XVE-SCL28* plants). **B.** DNA ploidy level distribution assessed by flow cytometry in nuclei isolated from the fourth leaf of 25 day-old wild-type (wt), *scl28-3* or *scl28-3 x ProSCL28:SCL28-VENUS* plants. **C.** DNA ploidy level distribution assessed by flow cytometry in nuclei isolated from whole roots of wild-type (wt), *scl28-3* and *scl28-3 x ProSCL28:SCL28-VENUS* plants. **D.** DNA ploidy level distribution in the root endodermis assessed by flow cytometry in wild-type (wt) and *scl28-3* plants expressing the *En7>>GFP* marker. **E.** DNA ploidy level distribution in root atrichoblasts assessed by flow cytometry in wild-type (wt) and *scl28-3* plants expressing the *GL2>>GFP* marker. **F.** Number of palisade parenchyma cells in mature leaves of wild-type (wt), *scl28-3* and *scl28-3 x ProSCL28:SCL28-VENUS* plants. Box plots with the measurements from 18 leaves are shown. Different letters indicate significant differences (*P* < 0.05; ANOVA followed by Tukey’s multiple comparison test). **G.** Number of cells counted in a single row of cortex from the QC to the junction between the root and the hypocotyl in 4-d-old wild-type (wt), *scl28-3* and *scl28-3* × *ProSCL28:SCL28-VENUS* plants. Different letters indicate significant differences (*P* < 0.05; ANOVA followed by Tukey’s multiple comparison test).

Endoreplication results in an increase in DNA content in the nuclei isolated from Arabidopsis roots and leaves. This results in a distribution of ploidy levels ranging from 2C to 32C in wild-type mature leaves (Figure 4B and Table S3A) and 2C to 16C in wild-type roots (Figure 4C and Table S3B). In *scl28-3* the distribution was shifted to lower ploidy levels. For example, in leaves, the percentage of nuclei with a DNA content of 32C was reduced from 9.2 in wild type to 1.3 in the mutant (Figure 4B and Table S3A). In whole roots, the reduction was less prominent, as the percentage of nuclei with a DNA content of 16C was reduced from 13.6 in wild type to 7.4 in the mutant (Figure 4C and Table S3B). However, it has been shown recently that each cell type has a particular endoreplication pattern (Bhosale et al., 2018). With this in mind, we measured in wild-type and *scl28-3* roots the DNA content of endodermis or atrichoblast cells only by flow cytometry using cell-type specific GFP marker lines (Table S4). For endodermis, a 10% shift from 8C to 4C was observed in *scl28-3* as compared to wild type (Figure 4D and Table S3C). In atrichoblasts, we observed in *scl28-3* a 3-fold decrease in the percentage of 16C cells with a concomitant increase in 8C and 4C cells (Figure 4E and Table S3D).

The observed changes in nuclear DNA content in whole leaves and roots were complemented by expressing *ProSCL28:SCL28-VENUS* (Figure 4B and C). Furthermore, overexpression of SCL28 in β-estradiol-treated *XVE-SCL28* roots caused a 6% increase in 16C cells at the expense of the lower ploidy levels (Figure S7 and Table S3E).

A defect in MCC exit and endoreplication onset might cause mitotic cells to persist for longer periods of time in that state and result in organs with an increased number of cells (Yi et al., 2014; Kumar et al., 2015). Indeed, the number of cells in mature leaves was increased by 25% in *scl28-3* as compared to wild type, a phenotype that was fully complemented by the *ProSCL28:SCL28-VENUS* construct (Figure 4F). Also, the number of cells present in single root cortex cell files of 4-d-old plants increased from 58 ± 4 in wild type to 76 ± 5 in the mutant and reverted to wild-type values with *ProSCL28:SCL28-VENUS* (Figure 4G).

Among the SCL28-regulated *SMRs*, *SMR13* (At5g59360) displayed substantial changes in response to changes in SCL28 expression (Figure 4A), so we focused on this gene for further analysis. A transcriptional reporter of *SMR13, ProSMR13:nlsGFP-GUS* (Yi et al., 2014), indicated that the CKI-coding gene is expressed in all tissues in a longitudinal gradient that starts in the transition zone (Figure 5A). When this reporter was introgressed into *scl28-3*, the detected GFP levels were reduced, resulting in a shootward displacement of the gradient and in lower plateau levels (Figure 5A and B). On the other hand, overexpression of *SCL28* in β-estradiol-treated *XVE-SCL28* plants increased the expression of the *ProSMR13:nlsGFP-GUS* reporter (Figure 5C and D). These results indicate that *SMR13* is expressed just before onset of endoreplication and is spatially and quantitatively regulated in the root meristem by *SCL28*. Finally, overexpression of *SMR13* in *scl28-3* plants (Figure S8) partially complemented the defects in both endoreplication (Figure 5E) and cell expansion (Figure 5F-L) in two independent transgenic lines transformed with a *Pro35S:SMR13* construct, supporting a role for *SCL28* in promoting endoreplication and cell expansion, presumably by activating *SMR13* expression.

**Figure 5.**
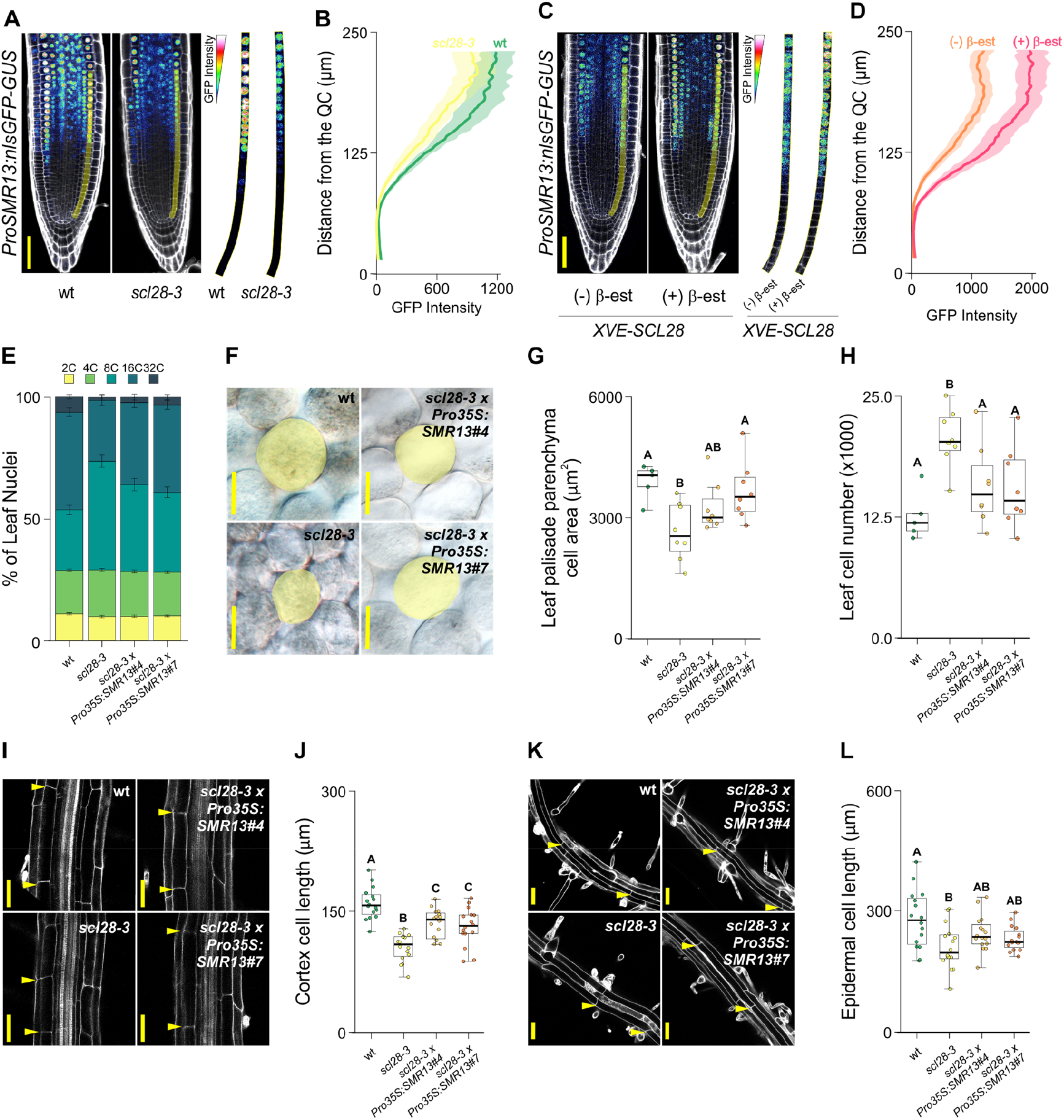
SCL28 promotes endoreplication and cell expansion by activating SMR13. **A.** Expression pattern of *SMR13* determined by LSCM using a *ProSMR13:nlsGFP-GUS* reporter (rainbow look up table) in wild-type (wt) and *scl28-3* roots. Images were obtained by LSCM in PI (grayscale) stained roots. Scale bars, 50 μm. To the right, a single row of cortex cells was segmented out for better visualization of the expression gradients. **B.** The expression gradient of *SMR13* in wild type and *scl28-3* was estimated by analyzing the fluorescence intensity profile in the root cortex of plants expressing the *ProSMR13:nlsGFP-GUS* reporter. The intensity profile corresponds to the average of 15 cortex cell files per genotype. **C.** Expression pattern of *SMR13* determined by LSCM using a *ProSMR13:nlsGFP-GUS* reporter (rainbow look up table) in plants over-expressing *SLC28*. Images were obtained by LSCM in PI (grayscale) stained *XVE-SCL28* plants grown in MS media supplemented with 0.25 μM β-estradiol. Scale bars, 50 μm. To the right, a single row of cortex cells was segmented out for better visualization of the expression gradient. **D.** The expression gradient of *SMR13* in plants over-expressing *SCL28* was estimated by analyzing the fluorescence intensity profile in the root cortex of β-estradiol-treated *XVE-SCL28* plants expressing the *ProSMR13:nlsGFP-GUS* reporter. The intensity profile corresponds to the average of 25 cortex cell files per condition. **E.** DNA ploidy level distribution assessed by flow cytometry in nuclei isolated from the fourth leaf of 25 day-old wild type (wt), *scl28-3* and two independent transgenic lines of *scl28-3* transformed with *Pro35S:SMR13*. **F.** View of cells from leaf palisade parenchyma in wild type (wt), *scl28-3* and two independent transgenic lines of *scl28-3* transformed with *Pro35S:SMR13*. Images were obtained by DIC microscopy in fixed and cleared leaves. Scale bars, 50 μm. The limits of representative cells are shaded in yellow. **G.** Leaf palisade parenchyma cell area in wild type (wt), *scl28-3* and two independent transgenic lines of *scl28-3* transformed with *Pro35S:SMR13*. Different letters indicate significant differences (*P* < 0.05; ANOVA followed by Tukey’s multiple comparison test). **H.** Number of palisade parenchyma cells in mature leaves of wild type (wt), *scl28-3* and two independent transgenic lines of *scl28-3* transformed with *Pro35S:SMR13*. Box plots with the measurements from 8 leaves are shown. Different letters indicate significant differences (*P* < 0.05; ANOVA followed by Tukey’s multiple comparison test). **I.** Mature root cortex cells in wild type (wt), *scl28-3* and two independent transgenic lines of *scl28-3* transformed with *Pro35S:SMR13*. Images were obtained by LSCM in PI (grayscale) stained roots. Scale bars, 50 μm. The limits of representative cells are labelled with arrow heads. **J.** Length of mature root cortex cells in wild type (wt), *scl28-3* and two independent transgenic lines of *scl28-3* transformed with *Pro35S:SMR13*. Box plots with the measurements of cells from 20 plants are shown. Different letters indicate significant differences (*P* < 0.05; ANOVA followed by Tukey’s multiple comparison test). **K.** Mature root epidermal cells in wild type (wt), *scl28-3* and two independent transgenic lines of *scl28-3* transformed with *Pro35S:SMR13*. Images were obtained by LSCM in PI (grayscale) stained roots. Scale bars, 50 μm. The limits of representative cells are labelled with arrow heads. **L.** Length of mature root epidermal cells in wild type (wt), *scl28-3* and two independent transgenic lines of *scl28-3* transformed with *Pro35S:SMR13*. Box plots with the measurements of cells from 20 plants are shown. Different letters indicate significant differences (*P* < 0.05; ANOVA followed by Tukey’s multiple comparison test).

## Discussion

We have previously shown that in Arabidopsis, *SCL28* stimulates plant organ growth by facilitating progression through G2/M in root meristems and modulates the selection of cell division planes, phragmoplast activity and mitotic cell expansion (Goldy et al., 2021). We now show that *SCL28* functions outside the cell proliferation domain promoting MCC exit and endoreplication onset by activating a group of CKIs that are expressed specifically in the transition and elongation zones (Yi et al., 2014).

We also found evidence that SCL28 modulates anisotropic post-mitotic cell expansion in several cell types from various plant organs, presumably by controlling the expression of genes involved in the biogenesis, assembly and remodeling of the cytoskeleton and cell wall. It remains to be determined whether this transcription factor regulates these cell wall, cytoskeletal and CKI genes directly or whether some genes are downstream to others. It has been proposed that SMR1-mediated endoreplication might exert an effect on cell growth through transcriptional control of cell wall genes that facilitate the cell wall changes required to support turgor-driven cell expansion (Bhosale et al., 2018; Bhosale et al., 2019). Related to this, a plausible hypothesis is that the changes in the expression of cell wall and cytoskeleton genes observed in *scl28-3* are a downstream consequence of the defects in endoreplication due to reduced *SMR* expression in the mutant.

We have previously shown that *SCL28* is activated by the R1R2R3-type MYB (MYB3R) MYB3R4 transcription factor and expressed at the highest levels in cells in the G2/M phases of the MCC (Goldy et al., 2021). MYB3R4 stimulates the expression of G2/M-specific genes in meristems through binding to a specific cis-acting regulatory sequence known as Mitotic Specific Activator (MSA) element (Kobayashi et al., 2015a). Thus, the expression of *SCL28* in the transition and elongation zones might result from the activity of other transcriptional regulators operating in these specific developmental zones or by the association of MYB3R transcription factors in higher order regulatory complexes (Kobayashi et al., 2015a; Kobayashi et al., 2015b).

Several transcription factors have been shown to operate in the transition zone to control meristem size, mitotic cell cycle and eventually, endoreplication onset. For example, *UPBEAT1* directly regulates the expression of a set of peroxidases that modulate the balance of reactive oxygen species and defines the size of the proliferation zone (Tsukagoshi et al., 2010) (Yamada et al., 2020). Also, *ARABIDOPSIS RESPONSE REGULATOR 1* (*ARR1*) regulates the expression of *EXPA1* in the transition zone. *EXPA1*, together with *EXPA10* and *EXPA15*, codes for a cell-wall located α-expansin that promotes cell expansion and mitotic cell cycle exit when activated by low apoplastic pH (Cosgrove, 2005). We note that *EXPA10* and *EXPA15* are not directly regulated by *ARR1* (Pacifici et al., 2018), but do respond to *SCL28* levels, suggesting that the GRAS-TF might promote mitotic cell cycle exit and cell expansion by activating a complementary group of expansins to those activated by *ARR1*.

Meristematic activity is regulated by external cues and in this way, plants re-configure their growth according to external conditions (Shimotohno et al., 2021). This is often accomplished by the induction of CKIs that drive MCC exit and eventually endoreplication. For example, the *SUPPRESSOR OF GAMMA RESPONSE 1* (*SOG1*) transcription factor activates the expression of *SMR5* and *SMR7* under various DNA damaging conditions (Yi et al., 2014; Bourbousse et al., 2018; Takahashi et al., 2019). In turn, induction of *SMR1* and *SMR5* under mild drought stress (Dubois et al., 2018) inhibit meristem activity restricting the growth of leaves and roots. As we found that a subset of *SMR* genes, most conspicuously *SMR9* and *SMR13* are regulated by *SCL28*, it remains to be determined if the GRAS TF also responds to environmental conditions and contributes to modulate the plasticity of plant development in response to environmental signals.

Our results add *SCL28* to the list of complex gene regulation mechanisms that occur in the transition zone. Such complexity of regulation is expected, considering the vast changes at the cellular level that occur in this area where cells transition from limited differentiation and active cell proliferation to cessation of division and enter their final program of expansion and differentiation. Thus, *SCL28* appears to modulate organ growth not only by facilitating progression through the phases of the cell cycle, but also by defining the size of the meristem by regulating MCC exit, endoreplication onset and cell expansion and differentiation, the latter probably indirectly by promoting an increase in somatic polyploidy.

## Methods

### Plant Materials and Growth Conditions

*Arabidopsis thaliana* accession Col-0 was used throughout this study. See Table S4 for a list and description of the mutants and reporter lines used in this study. Plants were grown in soil in long photoperiods (16 h light/8 h dark) using fluorescent bulbs (triphosphor code #840, 100 μmol quanta m^−2^ s^−1^) at 21°C. To perform root phenotype analysis, surface-sterilized seeds were sown on Petri plates containing Murashige and Skoog (MS) solid medium containing 1 × Murashige and Skoog salt mixture, 1% sucrose, and 2.3 mM MES (pH 5.8 adjust with KOH) in 1% agar. For the induction of the *XVE-SCL28* line, media was supplemented with 0.25 μM β-estradiol for the indicated times. For the experiments at pH4, the pH of the media was adjusted using HCl. Petri plates were placed in the dark at 4°C for 2 days. After stratification, Petri dishes were placed in a vertical orientation inside growth chambers in continuous light condition at 21 °C. See Table S5 for a detailed description of the constructs prepared and used in this study. Arabidopsis plants were transformed using the floral dip method (Clough and Bent, 1998).

### scl28-3 transcriptome analysis

Total RNA was isolated from root tips (~2 mm) in three biological replicates for each genotype using RNeasy plant mini kit (QIAGEN, http://www.qiagen.com). Approximately 300 root tips from 6-days-old plants were collected for each sample. The total RNA was subsequently used to prepare libraries for Illumina RNA-sequencing. For the analysis of gene expression, after removal of adaptors and quality assessment of the sequencing, reads were aligned to the TAIR10 release of the Arabidopsis genome (Lamesch et al., 2012) using hisat2 version 2.0.0-beta (Kim et al., 2019) with default parameters. Alignment data were sorted by Samtools version 1.7 (Danecek et al., 2021) and gene counts were generated using htseq-count version 0.6.1p1 (Anders et al., 2015) with parameter “--stranded=reverse.” Counts were processed in edgeR version 3.20.9 (Robinson et al., 2010), R version 3.4.4 (Team, 2020). Genes were considered expressed if counts were observed in 3 or more libraries. Wildtype and mutant samples were designated as respective treatment groups and statistical tests for difference in expression were performed using the glmQLFit and glmQLFTest functions. Analysis scripts are available upon request. Genes with an unadjusted p-value < 0.05 and fold change > 25% were considered as differentially expressed and chosen for further study and validation.

GO terms enrichment in the lists of differentially expressed genes were obtained with the GO-TermFinder (Boyle et al., 2004) via the Princeton GO-TermFinder web server (https://go.princeton.edu/cgi-bin/GOTermFinder) and simplified using REVIGO (Supek et al., 2011).

### Gene Expression Analysis via RT-qPCR

Total RNA was isolated from roots using RNeasy plant mini kit (QIAGEN, http://www.qiagen.com). Approximately 150 root tips (~2 mm) from 6-days-old plants were collected for each sample. The numbers of biological replicates are specified in the corresponding figure legends. Total RNA (0.5 μg) was treated with RQ1 RNase-free DNase (Promega). First-strand cDNA synthesis was performed using Moloney Murine Leukemia Virus Reverse Transcriptase (MMLV, Invitrogen). PCR was performed in an AriaMx thermal cycler (Agilent) using EvaGreen to monitor double-stranded DNA synthesis. Normalized relative quantities (NRQ) were obtained using the qBase method (Hellemans et al., 2007) with *RPS26C* and *PP2A* as reference genes for normalization across samples. When indicated, NRQ values were normalized to the mean value obtained in wild-type plants or control treatments. Melting curve analyses at the end of the process and “no template controls” were performed to ensure product-specific amplification without primer-dimer artifacts. Primer sequences are given in Table S6.

### Confocal Microscopy

Laser scanning confocal microscopy (LSCM) was performed throughout the study using a Plan Apochromat 20×, 0.8-NA lens on a Zeiss LSM880 microscope. Roots, hypocotyls, and stems were stained with 15 μg/mL Propidium Iodide (PI) (Sigma) for 3 min (roots) or 30 min (hypocotyls and stems). Organs were mounted in water and visualized after excitation by a 488-nm laser line for GFP, PI or by a 514-nm laser line for VENUS. The fluorescence emission was collected between 590 and 700 nm for PI, 496 and 542 nm for GFP and 524 and 570 nm for VENUS. Cellular parameters and fluorescence signal intensity were analyzed with using FIJI (Schindelin et al., 2012). Fluorescence intensity measurements in *ProSMR13-nlsGUS:GFP* lines were performed as described (Ercoli et al., 2018).

### Analysis of phenotypes

For root-elongation measurements, seedlings were grown vertically for the indicated number of days. Starting from day 5 after germination until the end of the experiment, a dot was drawn at the position of the root tip. Finally, plates were photographed, and the root length was measured over time. Root growth rate, expressed in millimeters per hour, was estimated from root length (millimeters) vs. plant age (days after sowing) plots. For rosette growth measurements, plants were grown in soil as described above a photographed daily from day 12 to 20 after sawing. The rosette area was measured using FIJI (Schindelin et al., 2012) and growth rate, expressed in square millimeters per hour, was estimated from rosette area (square millimeters) vs. plant age (days after sowing) plots. For inflorescence stem growth measurements, plants were grown in soil and the length of inflorescence stems was measured daily from day 35 to 42 after sawing.

In roots, meristematic zone length and meristematic cell number were determined from LSCM images using the file of cortex cells in 6-days seedlings. The meristematic zone was defined as the region from the quiescent centre up to the cell that was twice the length of the immediately preceding cell (Dello Ioio et al., 2008). The length of mature root cortex or epidermis cells was measured in cells in the mature zone, where the epidermis hairs are already fully developed.

To obtain paradermal views of palisade parenchyma or leaf epidermal cells, leaves were fixed overnight with 70% ethanol and then cleared with lactic acid for at least 2 days (Fiorani and Beemster, 2006). Cells were observed using differential interference contrast microscopy (DIC) in an Olympus BH2 microscope. The area of at least 20 cells per leaf were measured using FIJI. The number of cells per organ was estimated by dividing the area of the leaf by the area of the corresponding cell type. Measurements were carried out in at least ten leaves.

The InfoStat software (https://www.infostat.com.ar/) was used to perform statistical analyses. The tests applied in each experiment are indicated in the legend of each figure.

### Ploidy analysis

For DNA content analysis in nuclei isolated from whole organs, leaves or roots were harvested and chopped with a razor blade in ice cold Galbraith`s extraction buffer (45 mM MgCl2; 30 mM sodium citrate; 20 mM 3-[N-morpholino] propanesulfonic acid, pH 7,0 and 0,1% Triton X-100). The suspension was filtered through a 20 μm nylon mesh to remove tissue debris. RNA was degraded adding a DNase-free RNase and nuclei were stained with 50 μg/ml PI. Finally, flow cytometry was performed in a BD FACSAria II cell sorter. The DNA content was derived from IP fluorescence measurements using the FlowJo software. These experiments were analyzed at least in triplicate and 3000 to 5000 stained nuclei were analyzed for each sample.

For flow cytometry on nuclear GFP lines, whole root from 6 days-old plants were chopped with a razor blade in 200 mL of Galbraith`s extraction buffer for 2 min, then filtered through a 50-mmnylon filter. The DNA was stained with 1 mg/mL DAPI (Zhang et al., 2005). Nuclei were measured using a Quantum P (Quantum Analysis) flow cytometer excited by illumination at 395 nm and equipped with an additional 488-nm laser to excite and detect GFP-specific fluorescence. The DNA content of GFP-positive nuclei was derived from DAPI fluorescence measurements using CyPAD software (Quantum Analysis).

## Acknowledgements

We thank members of the lab for comments and discussions. We thank the technical support from Diego Aguirre for plant work, Enrique Morales for microscopy and Mara J. Ojeda and Ilse Vercauteren for flow cytometry. CG, VB and CB were supported by fellowships from CONICET and ANPCyT. RER is members of CONICET. RER was supported by the Argentinean Ministry of Science (PICT2015-3758 and PICT2017-2762).

