## Supplementary Figures for "SCL28 promotes cell expansion and endoreplication in Arabidopsis by activating *SIAMESE-RELATED* cyclin-dependent kinase inhibitors"

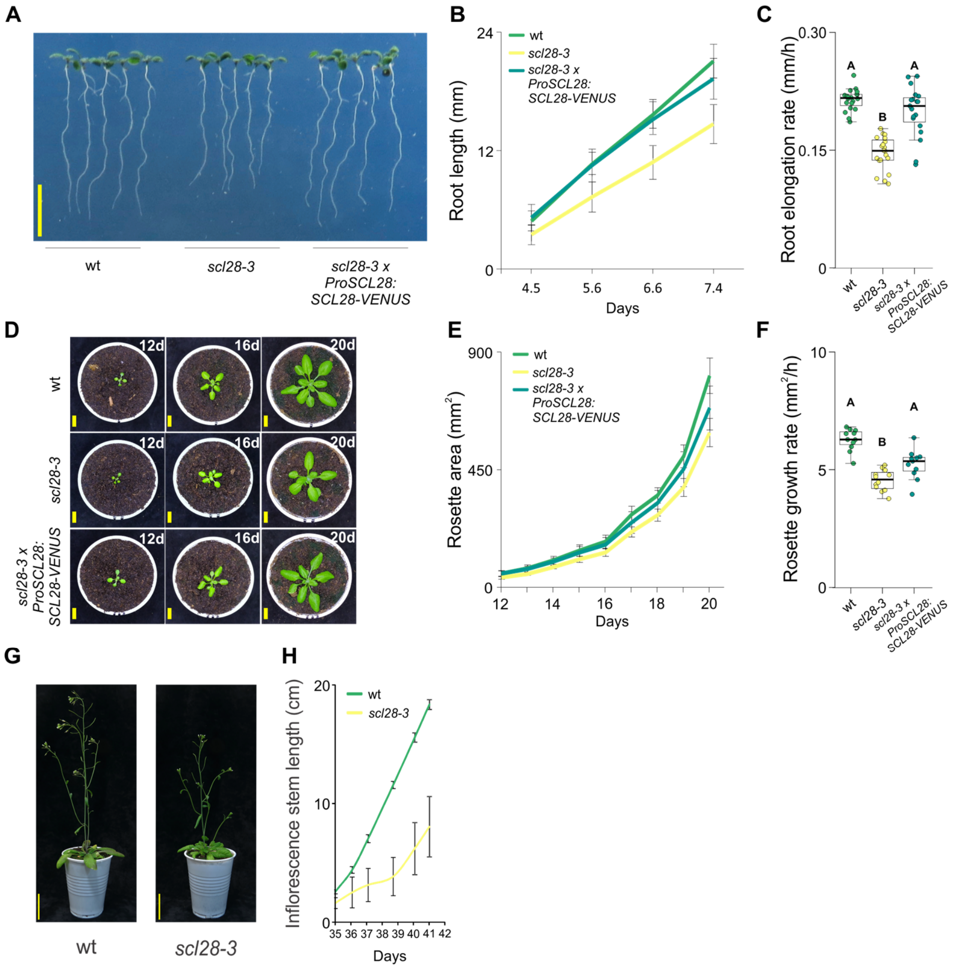


***Figure S1. SCL28 promotes growth of various organs***

A. Root phenotype of 10 d-old wild-type (wt), *scl28-3*, and *scl28-3×ProSCL28:SCL28-VENUS* plants. Scale bar, 1 cm.

B. Root growth of wild-type (wt), *scl28-3*, and *scl28-3×ProSCL28:SCL28-VENUS* plants.

C. Root elongation rate (mm/h) of wild-type, *scl28-3*, and *scl28-3×ProSCL28:SCL28-VENUS* plants. Box plots with the measurements from 20 roots are shown. Different letters indicate significant differences ($P < 0.05$; ANOVA followed by Tukey's multiple comparison test).

D. Rosette phenotype of 12, 16 and 24 d-old wild-type (wt), *scl28-3*, and *scl28-3×ProSCL28:SCL28-VENUS* plants. Scale bars, 1 cm.

E. Rosette growth of wild-type (wt), *scl28-3*, and *scl28-3×ProSCL28:SCL28-VENUS* plants.

F. Rosette growth rate ($mm^2$/h) of wild-type, *scl28-3*, and *scl28-3×ProSCL28:SCL28-VENUS* plants. Box plots with the measurements from 10 rosette are shown. Different letters indicate significant differences ($P < 0.05$; ANOVA followed by Tukey's multiple comparison test).

G. Shoot phenotype in 35 d-old wild-type (wt) and *scl28-3* plants. Scale bar, 1 cm.

H. Inflorescence stem growth of wild-type (wt) and *scl28-3* plants.

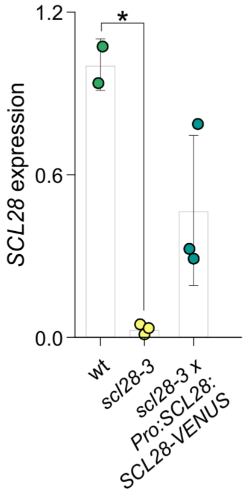


***Figure S2. SCL28 expression in wt, scl28-3 and scl28-3 x ProSCL28:SCL28-VENUS plants***
Expression of *SCL28* in wild-type (wt), *scl28-3* and *scl28-3* x *ProSCL28:SCL28-VENUS* plants. Expression was estimated by RT-qPCR in three biological replicates and normalized to the mean value obtained in wild-type plants. Asterisks indicate significant differences (Student's t test, $p<0.05$).

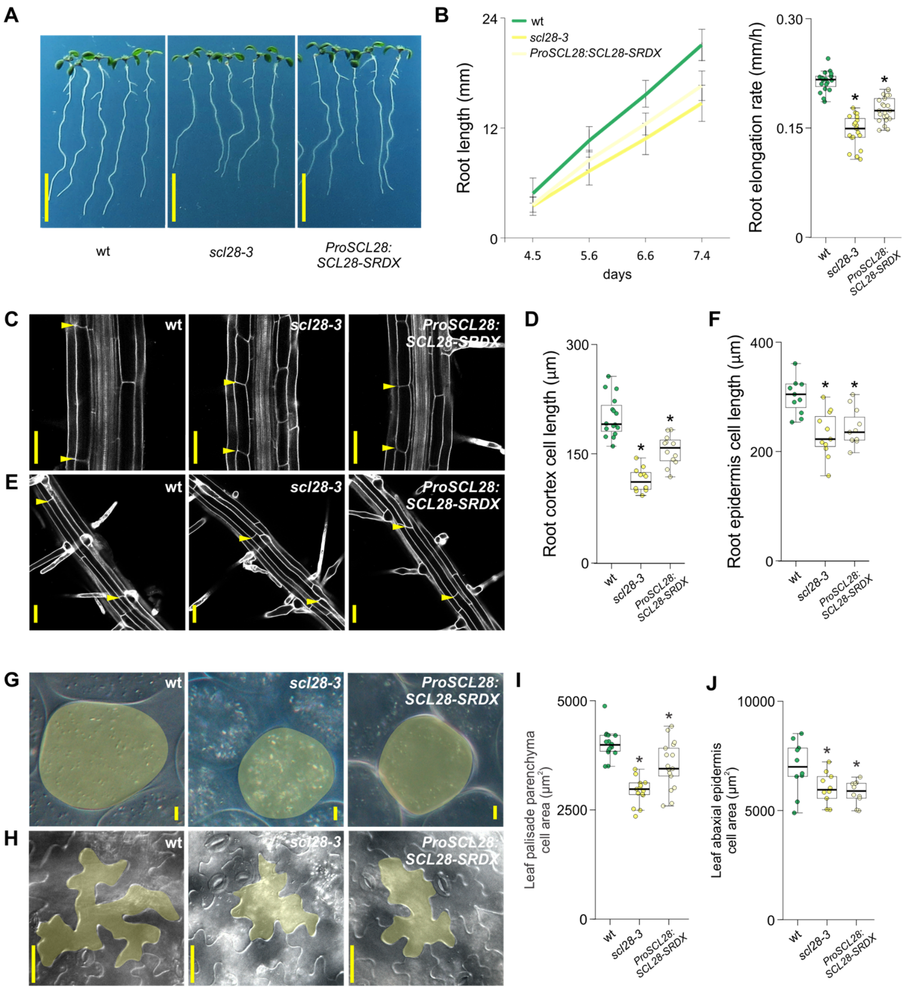


***Figure S3. Plants expressing SCL28-SRDX phenocopiesthe scl28-3 mutant***

**A.** Root phenotype of 10 d-old wild-type (wt), *scl28-3*, and *ProSCL28:SCL28-SRDX* plants. Scale bar, 1 cm.

**B.** Root growth of wild-type (wt), *scl28-3*, and *ProSCL28:SCL28-SRDX* plants (right) and root elongation rate (left) of wild-type, *scl28-3*, and *ProSCL28:SCL28-SRDX* plants. Box plots with the measurements from 20 roots are shown. Asterisks indicate significant differences with wild type ($P < 0.05$, Student's t test).

**C**. Mature root cortex cells in wild-type (wt), *scl28-3* and *ProSCL28:SCL28-SRDX* plants. Images were obtained by LSCM in PI (grayscale) stained roots. Scale bars, 50 µm.

**D**. Length of mature root cortex cells in wild-type (wt), *scl28-3* and *ProSCL28:SCL28-SRDX* plants. Box plots with the measurements of cells from 20 plants are shown. Asterisks indicate significant differences with wild type ($P < 0.05$, Student's t test).

**E**. Mature root epidermal cells in wild-type (wt), *scl28-3* and *ProSCL28:SCL28-SRDX* plants. Images were obtained by LSCM in PI (grayscale) stained roots. Scale bars, 50 µm.

**F**. Length of mature root epidermal cells in wild-type (wt), *scl28-3* and *ProSCL28:SCL28-SRDX* plants. Box plots with the measurements of cells from 20 plants are shown. Asterisks indicate significant differences with wild type ($P < 0.05$, Student's t test).

**G-H.** View of cells from leaf palisade parenchyma (G) and leaf abaxial epidermis (H) in wild-type (wt), *scl28-3* and *ProSCL28:SCL28-SRDX* plants. Images were obtained by DIC microscopy in fixed and cleared leaves. Representative cells are shaded in yellow. Scale bars, 50 μm.

**I-J.** Cell area measurements of leaf palisade parenchyma (I) and leaf abaxial epidermis (I) cells in wild-type (wt), *scl28-3* and *ProSCL28:SCL28-SRDX* plants. Box plots with the measurements of cells from 20 plants are shown. Asterisks indicate significant differences with wild type ($P < 0.05$, Student's t test).

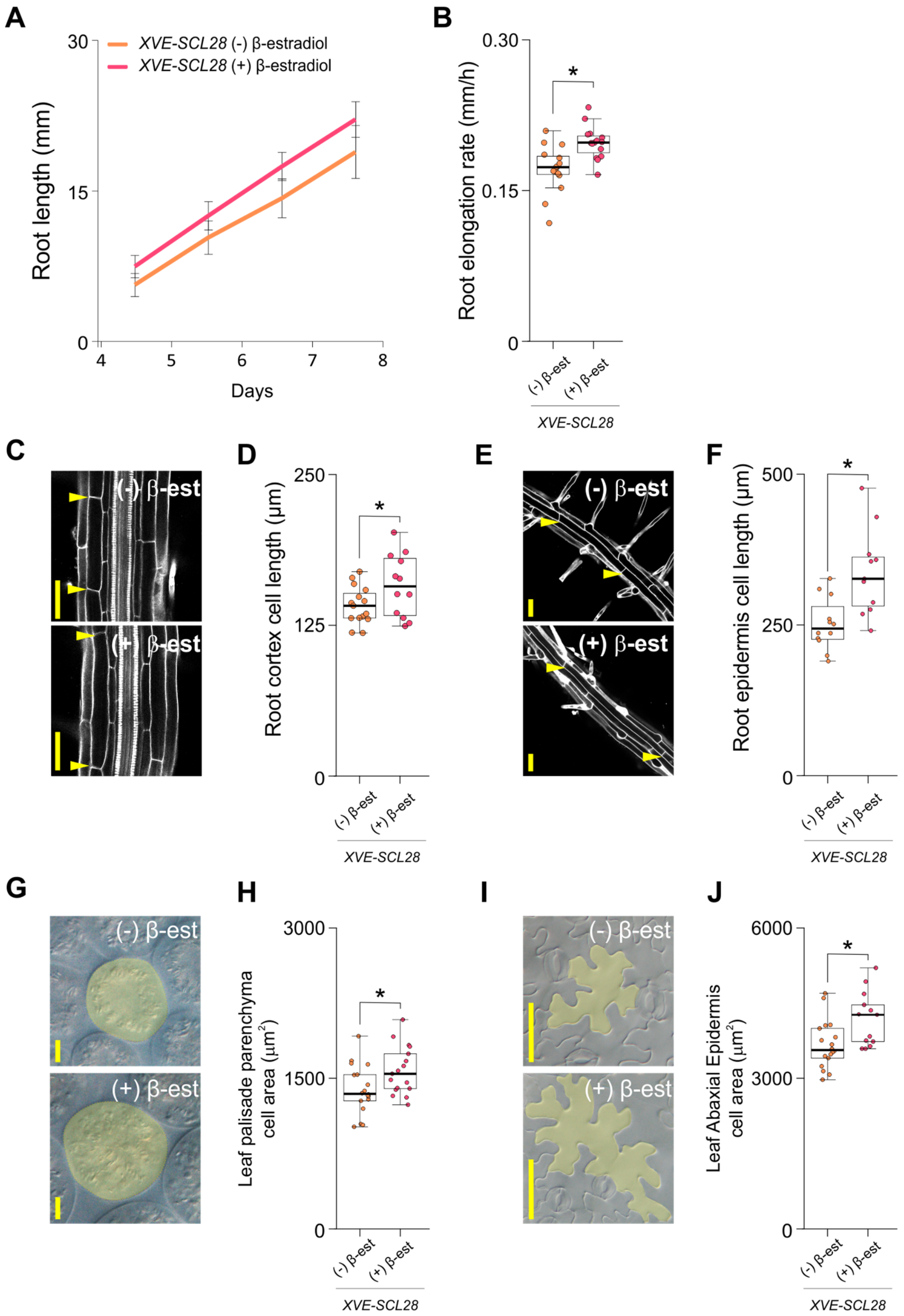


***Figure S4. Overexpression of SCL28 in β-estradiol-treated XVE-SCL28 plants leads to phenotypes opposite to those observed in scl28-3***

**A.** Root growth of *XVE-SCL28* plants grown in 0.25 µM β-estradiol.

**B.** Root elongation rate of*XVE-SCL28* plants grown in β-estradiol. Box plots with the measurements from 20 roots are shown. Asterisks indicate significant differences with mock-treated plants (*P* < 0.05, Student's t test).

**C**. Mature root cortex cells from *XVE-SCL28* plants grown in β-estradiol. Images were obtained by LSCM in PI (grayscale) stained roots. Scale bars, 50 μm. The limits of representative cells are labelled with arrowheads.

**D**. Length of mature root cortex cells from *XVE-SCL28* plants grown in β-estradiol. Box plots with the measurements of cells from 20 plants are shown. Asterisks indicate significant differences with mock-treated plants ($P < 0.05$, Student's t test).

**E**. Mature root epidermal cells of *XVE-SCL28* plants grown in β-estradiol. Images were obtained by LSCM in PI (grayscale) stained roots. Scale bars, 50 μm. The limits of representative cells are labelled with arrowheads.

**F**. Length of mature root epidermal cells of *XVE-SCL28* plants grown in β-estradiol. Box plots with the measurements of cells from 20 plants are shown. Asterisks indicate significant differences with mock-treated plants ($P < 0.05$, Student's t test).

**G-H.** View leaf palisade parenchyma (G) and leaf abaxial epidermis (H) cells from *XVE-SCL28* plants grown in β-estradiol. Images were obtained by DIC microscopy in fixed and cleared leaves. Representative cells are shaded in yellow. Scale bars, 50 μm.

**I-J.** Area of leaf palisade parenchyma (I) and leaf abaxial epidermis (I) cells from *XVE-SCL28* plants grown in β-estradiol. Box plots with the measurements of cells from 20 plants are shown. Asterisks indicate significant differences with mock-treated plants ($P < 0.05$, Student's t test).

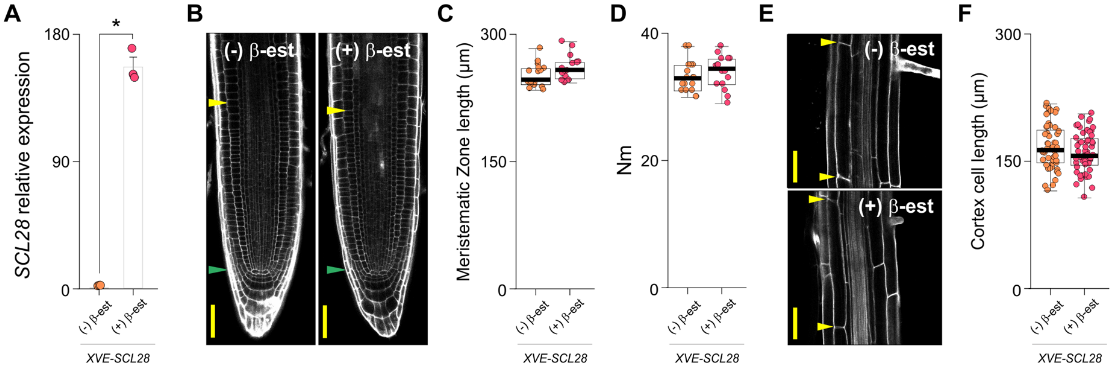


***Figure S5. Expression of SCL28 and absence of phenotypes in XVE-SCL28 plants treated with β-estradiol for short periods***

A. Expression of *SCL28* in *XVE-SCL28* plants transplanted to MS media supplemented with 0.25 µM β-estradiol for 16 h. Expression was estimated by RT-qPCR in three biological replicates and normalized to the mean value obtained in mock-treated control plants. Asterisks indicate significant differences ($P < 0.05$, Student's t test).

B-D. Root tip architecture (B), meristematic zone length (C) and number of meristematic cortex cells (Nm, D) of 6-d-old *XVE-SCL28* plants transplanted to MS media supplemented with 0.25 µM β-estradiol for 16 h. In B, green and yellow arrowheads mark the position of the QC and the end of the meristem where cells start to elongate. Scale bars, 50 µm.

E. Mature cortex cells in *XVE-SCL28* plants transplanted to MS media supplemented with 0.25 µM β-estradiol for 16 h. Images were obtained by LSCM in PI (grayscale) stained organs. The limits of representative cells are labelled with arrowheads. Scale bars, 50 µm.

F. Length of mature cortex cells in *XVE-SCL28* plants transplanted to MS media supplemented with 0.25 µM β-estradiol for 16 h. Box plots with the measurements of 50 cells are shown.

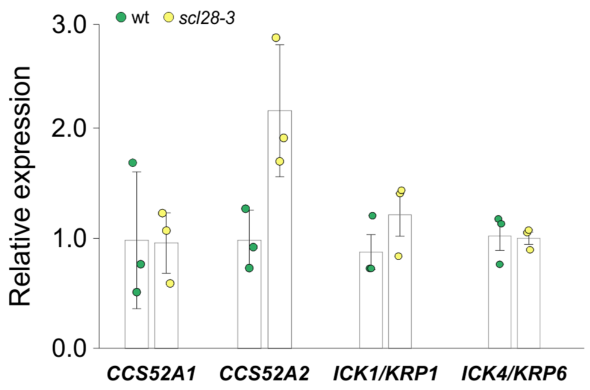


***Figure S6. Expression of cell-cycle regulators in plants with altered levels of SCL28***
Expression of *CCS52A1*, *CCS52A2*, *ICK1/KRP1*, *ICK4/KRP6,* in wild type (wt) and *scl28-3*. Expression was estimated by RT-qPCR in three biological replicates and normalized to the mean value obtained in wild-type plants. Asterisks indicate significant differences ($P < 0.05$, Student's t test).

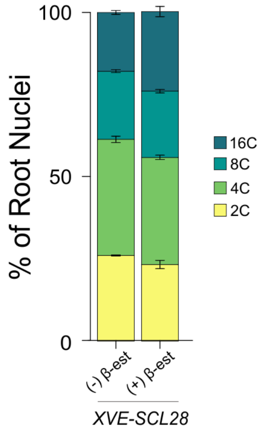


***Figure S7. SCL28 overexpression enhances endoreplication***
DNA ploidy level distribution assessed by flow cytometry in nuclei isolated from whole roots of *XVE-SCL28* plants grown in MS media supplemented with 0.25 µM β-estradiol.

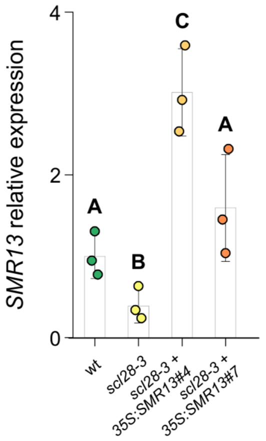


***Figure S8. Expression of SMR13 in scl28-3 plants transformed with Pro35S:SMR13***

Expression of *SMR13* in wild type (wt), *scl28-3* and two independent transgenic lines of *scl28-3* transformed with *Pro35S:SMR13*. Expression was estimated by RT-qPCR in three biological replicates and normalized to the mean value obtained in wild-type plants. Different letters indicate significant differences ($P < 0.05$; ANOVA followed by Tukey's multiple comparison test).
