## Supplementary Tables for "SCL28 promotes cell expansion and endoreplication in Arabidopsis by activating *SIAMESE-RELATED* cyclin-dependent kinase inhibitors"

**Table S1. SCL28 promotes cell expansion in multiple plant organs.**
Cell size in various cell types from roots and shoots in wild-type (wt), *scl28-3* and *scl28-3* × *ProSCL28:SCL28-VENUS* plants.

**Table S2. Transcriptome analysis of *scl28-3*.**
Table S2A. Expression of detected genes in the transcriptome analysis is informed as fold change (*scl28-3*/wild-type) together with the corresponding unadjusted p value for the comparison across genotypes.
Table S2B. Fold change (*scl28-3*/wild-type) and unadjusted p value for genes up-regulated in scl28-3. The name of selected genes further studied in this work is indicated.
Table S2C. Fold change (*scl28-3*/wild-type) and unadjusted p value for genes down-regulated in scl28-3. The name of selected genes further studied in this work is indicated.
Table S2D. GO term enrichment analysis in the list of differentially expressed genes.

**Table S3 . DNA ploidy level distribution.**
Table S3A. DNA ploidy level distribution in leaves from wild type (wt), *scl28-3* and *scl28-3* x *ProSCL28:SCL28-VENUS*.
Table S3B. DNA ploidy level distribution in roots from wild type (wt), *scl28-3* and *scl28-3* x *ProSCL28:SCL28-VENUS*.
Table S3C. DNA ploidy level distribution in nuclei isolated from root endodermis cells from wild-type (wt) and *scl28-3* plants.
Table S3D. DNA ploidy level distribution in nuclei isolated from root atrichoblast cells from wild-type (wt) and *scl28-3* plants.
Table S3E. DNA ploidy level distribution in roots from plants over-expressing *SCL28*.
Table S3F. DNA ploidy level distribution in leaves from wild type (wt), *scl28-3* and *scl28-3* x *Pro35S:SMR13*.

**Table S4. Mutants and reporter lines used in this study.**

**Table S5. Binary plasmids generated in this study.**

**Table S6. Oligonucleotide primers used in this study.**

**Table S3. DNA ploidy level distribution.**

*Table S3A. DNA ploidy level distribution in leaves from wt, scl28-3 and scl28-3 x ProSCL28:SCL28-VENUS.*

| Ploidy (%) | wt | *scl28-3* | *scl28-3 x ProSCL28:SCL28--VENUS* |
|---|---|---|---|
| *2C* | 10.8 ± 0.5$^{A}$ | 11.2 ± 0.2$^{A}$ | 8.8 ± 0.5$^{B}$ |
| *4C* | 16.3 ± 0.5$^{A}$ | 17.6 ± 0.2$^{A}$ | 16.2 ± 0.5$^{A}$ |
| *8C* | 19.6 ± 0.2$^{A}$ | 38.3 ± 0.6$^{B}$ | 17.9 ± 1.0$^{A}$ |
| *16C* | 43.6 ± 1.4$^{A}$ | 31.2 ± 0.8$^{B}$ | 41.4 ± 1.3$^{A}$ |
| *32C* | 9.2 ± 1.0$^{A}$ | 1.3 ± 0.1$^{B}$ | 15.4 ± 1.7$^{C}$ |

$^{A,B}$ Within each ploidy level, different letters indicate significant differences (P < 0.05; ANOVA followed by Tukey's multiple comparison test).

*Table S3B. DNA ploidy level distribution in roots from wt, scl28-3 and scl28-3 x ProSCL28:SCL28-VENUS.*

| Ploidy (%) | wt | *scl28-3* | *scl28-3 x ProSCL28:SCL28--VENUS* |
|---|---|---|---|
| *2C* | 35.4 ± 1.5$^{A}$ | 41.0 ± 2.1$^{B}$ | 37.1 ± 0.8$^{AB}$ |
| *4C* | 32.0 ± 0.3$^{A}$ | 31.7 ± 1.0$^{A}$ | 31.0 ± 0.5$^{A}$ |
| *8C* | 19.1 ± 0.7$^{A}$ | 20.0 ± 1.7$^{A}$ | 16.4 ± 0.5$^{B}$ |
| *16C* | 13.6 ± 1.0$^{A}$ | 7.4 ± 0.3$^{B}$ | 15.6 ± 0.9$^{A}$ |

$^{A,B}$ Within each ploidy level, different letters indicate significant differences (P < 0.05; ANOVA followed by Tukey's multiple comparison test).

*Table S3C. DNA ploidy level distribution in nuclei isolated from root endodermis cells from wt and scl28-3 plants.*

| Ploidy (%) | wt | *scl28-3* |
|---|---|---|
| *2C* | 20.7 ± 1.2 | 20.4 ± 0.5 |
| *4C* | 48.5 ± 1.0 | 58.6 ± 0.7* |
| *8C* | 28.5 ± 1.6 | 18.4 ± 0.6* |
| *16C* | 2.3 ± 0.2 | 2.6 ± 0.3 |

$^{A,B}$ Within each ploidy level, asterisks indicate significant differences (P < 0.05; Student´s t test).

*Table S3D. DNA ploidy level distribution in nuclei isolated from root atrichoblast cells from wt and scl28-3 plants.*

| Ploidy (%) | wt | *scl28-3* |
|---|---|---|
| *2C* | 8.4 ± 1.0 | 7.8 ± 0.3 |
| *4C* | 7.9 ± 0.7 | 9.8 ± 0.4* |
| *8C* | 73.3 ± 1.1 | 78.9 ± 0.7* |
| *16C* | 10.4 ± 0.4 | 3.6 ± 0.3* |

$^{A,B}$ Within each ploidy level, asterisks indicate significant differences (P < 0.05; Student´s t test).

*Table S3E. DNA ploidy level distribution in roots from plants over-expressing SCL28.*

| Ploidy (%) | *XVE-SCL28* | *XVE-SCL28* + β-estradiol |
|---|---|---|
| *2C* | 26.0 ± 1.2 | 23.8 ± 0.2 |
| *4C* | 35.2 ± 0.7 | 32.5 ± 1.0 |
| *8C* | 20.7 ± 0.5 | 20.1 ± 0.5 |
| *16C* | 17.8 ± 1.5 | 24.1 ± 0.6* |

*Within each ploidy level, asterisks indicate significant differences (P < 0.05; Student´s t test).

*Table S3F. DNA ploidy level distribution in leaves from wild-type, scl28-3 and scl28-3 x Pro35S:SMR13.*

| Ploidy (%) | wt | *scl28-3* | *scl28-3* x *Pro35S:SMR13* #4 | *scl28-3* x *Pro35S:SMR13* #8 |
|---|---|---|---|---|
| *2C* | 11.0 ± 0.4$^{A}$ | 9.9 ± 0.5 $^{A}$ | 10.0 ± 0.5 $^{A}$ | 10.2 ± 0.4 $^{A}$ |
| *4C* | 17.9 ± 0.5 $^{A}$ | 19.2 ± 0.6 $^{A}$ | 18.5 ± 0.6 $^{A}$ | 17.9 ± 0.5 $^{A}$ |
| *8C* | 24.8 ± 1.9 $^{A}$ | 44.6 ± 2.5 $^{C}$ | 35.6 ± 2.5 $^{B,C}$ | 32.8 ± 2.2 $^{A,B}$ |
| *16C* | 40.1 ± 1.7 $^{A}$ | 25.0 ± 2.1 $^{B}$ | 33.7 ± 2.1 $^{A}$ | 35.9 ± 1.8 $^{A}$ |
| *32C* | 6.3 ± 0.9 $^{A}$ | 1.2 ± 1.2 $^{B}$ | 2.2 ± 1.2 $^{A,B}$ | 3.2 ± 1.0 $^{A,B}$ |

$^{A,B,C}$ Within each ploidy level, different letters indicate significant differences (P < 0.05; ANOVA followed by Tukey's multiple comparison test).

**Table S4. Mutants and reporter lines used in this study.**

| Line | Line Description | Reference |
|---|---|---|
| ***scl28-3*** | Insertional mutant. SALK_205284 | Goldy *et al*. (2021) *PNAS* |
| ***ProSCL28:SCL28-VENUS*** | SCL28 Reporter | Goldy *et al*. (2021) *PNAS* |
| ***ProSCL28:SCL28-SRDX*** | SCL28-derived transcriptional repressor | Goldy *et al*. (2021) *PNAS* |
| ***XVE-SCL28*** | β-estradiol *SCL28* inducible line | Coego *et al.* (2014) The Plant Journal |
| ***ProSMR13-nlsGUS:GFP*** | *SMR13* transcriptional marker | Yi *et al*. (2014) The Plant Cell |
| ***GL2>>GFP*** | Root epidermis atrichoblast marker | Dietrich *et al.* (2017) Nature Plants |
| ***En7>>GFP*** | Root endodermis marker | Dietrich *et al.* (2017) Nature Plants |

**Table S5. Binary plasmids generated in this study.**

| Vector | Construct | Arabidopsis Chromosome: Start-End |
|---|---|---|
| **CG28** | *Pro35S:SMR13-GFP* | *Pro35S* : [*SMR13* CDS, 5: 23946059 - 23945595] – GFP |

All constructs were cloned in the binary vector pCHF3 (Jarvis, P., Chen, L. J., Li, H., Peto, C. A., Fankhauser, C., and Chory, J. (1998). An Arabidopsis mutant defective in the plastid general protein import apparatus. Science 282, 100103). T-DNA constructs were introduced into *A. tumefaciens* strain ASE (Fraley, R. T., Rogers, S. G., Horsch, R. B., Eichholtz, D. A., Flick, J. S., Fink, C. L., Hoffmann, N. L., and Sanders, P. R. (1985) The SEV system: a new disarmed Ti plasmid vector system for plant transformation. Biotechnology 3, 629635).

**Table S6. Oligonucleotide primers used in this study.**

| Gene | Locus ID | Sequence | Porpouse |
|---|---|---|---|
| ***SCL28*** | AT1G63100 | GATGAAGACAACGGCGGTGAAG | RT-qPCR Forward primer |
| | | TTCCCTCCGGTAGTCCAAGC | RT-qPCR Reverse primer |
| ***ADF6*** | AT2G31200 | TGTTGATGAGCATGATGAGAGA | RT-qPCR Forward primer |
| | | TGTTCGCAAGGATTAGACTCG | RT-qPCR Reverse primer |
| ***IQD16*** | AT4G10640 | AAAAACCGCCGGTGATTGTC | RT-qPCR Forward primer |
| | | ATAATGGCAGCCCAATGACG | RT-qPCR Reverse primer |
| ***EXT37*** | AT4G16140 | TCTCAATCTGGCTCTTACCGTCCA | RT-qPCR Forward primer |
| | | TGCGGCGGAGGGTTGTAATAGTAA | RT-qPCR Reverse primer |
| ***EXPA10*** | AT1G26770 | GATAAACGGCCACTCATACTTC | RT-qPCR Forward primer |
| | | GTGGTGACCTTAAAGGAAAGTG | RT-qPCR Reverse primer |
| ***EXPA15*** | AT2G03090 | TTGGTGTAACCCTCCGCTTCATCA | RT-qPCR Forward primer |
| | | GCAACCGAATGAACATCTCCAGCA | RT-qPCR Reverse primer |
| ***MAN7*** | AT5G66460 | AGCAGGAGGATTGTTCTGGCAACT | RT-qPCR Forward primer |
| | | TCAACTTCCGCGATTGCTGTGA | RT-qPCR Reverse primer |
| ***GH9C2*** | AT1G64390 | TGCTCCTGTTGTCCGAAGACCAAT | RT-qPCR Forward primer |
| | | AAACCGAACGAGTTGCCTGATCTC | RT-qPCR Reverse primer |
| **AT4G26830** | AT4G26830 | CGGTAACAGAGGTAAAGTTCCGGTGA | RT-qPCR Forward primer |
| | | ATTGAAGGCGTACGAAGCGTGA | RT-qPCR Reverse primer |
| ***XTH5*** | AT5G13870 | TAAGTTCTGCGAGACGCAGGGAAA | RT-qPCR Forward primer |
| | | ACGCGGTCAGTGCAATAGTTGT | RT-qPCR Reverse primer |
| ***AGP30*** | AT2G33790 | TCAACAACGTCCAAGGCGCTAA | RT-qPCR Forward primer |
| | | AGACTTCACGAGGAAAGCTCGACA | RT-qPCR Reverse primer |
| ***SIM*** | AT5G04470 | CACAAGATTCCTCCCACCACAG | RT-qPCR Forward primer |
| | | CAGAGGAGAAGAACCGCTCGAT | RT-qPCR Reverse primer |
| ***SMR1*** | AT3G10525 | CACCCACATCCCAAGAACACAAG | RT-qPCR Forward primer |
| | | GACGGAGGAGAAGAAACGGTCAA | RT-qPCR Reverse primer |
| ***SMR2*** | AT1G08180 | AGAGCAGAAACCCAGAAGCCAAG | RT-qPCR Forward primer |
| | | GAAATCTCACGCGGTCGCTTTCTT | RT-qPCR Reverse primer |
| ***SMR6*** | AT5G40460 | GGGCTTCGTTGAAACCAGTCAAG | RT-qPCR Forward primer |
| | | TTTCTCGGTGCTGGTGGACATTC | RT-qPCR Reverse primer |
| ***SMR7*** | AT3G27630 | GCCAAAACATCGATTCGGGCTTC | RT-qPCR Forward primer |
| | | TCGCCGTGGGAGTGATACAAAT | RT-qPCR Reverse primer |
| ***SMR9*** | AT1G51355 | GCCACTTCAAGAACCCATCTCC | RT-qPCR Forward primer |
| | | TCCGGAGTACAACATCCACTCTCT | RT-qPCR Reverse primer |
| ***SMR13*** | AT5G59360 | GAACCACCAACACCGACAACAAG | RT-qPCR Forward primer |
| | | GTTCGAGTTTCTCGGCGTCTCT | RT-qPCR Reverse primer |
| | | AATCGCGTAATGAACCCATCCACG | RT-qPCR Reverse primer |
| ***ICK1/KRP1*** | AT2G23430 | GGTGACACTGAAACGTCGAC | RT-qPCR Forward primer |
| | | CTCCGTCATCAATTTCGCCT | RT-qPCR Reverse primer |
| ***ICK4/KRP6*** | AT3G19150 | AGCTTCAAGCACAAGCTTCTCACC | RT-qPCR Forward primer |
| | | TTCTCCACCGATGGTAACGGAACA | RT-qPCR Reverse primer |
| ***RPS26C*** | AT3G56340 | GACTTTCAAGCGCAGGAATGGTG | RT-qPCR Forward primer |
| | | CCTTGTCCTTGGGGCAACACTTT | RT-qPCR Reverse primer |
| ***PP2A*** | AT1G13320 | CCTGCGGTAATAACTGCATCT | RT-qPCR Forward primer |
| | | CTTCACTTAGCTCCACCAAGCA | RT-qPCR Reverse primer |

[a] To genotype for the T-DNA insertions, the LBb1 primer (GCGTGGACCGCTTGCTGCAACT) was used.
